# Priority effects among amphibian microbes lead to varying levels of Bd infection

**DOI:** 10.1101/2021.07.20.453099

**Authors:** Elle M. Barnes, J.D. Lewis

**Affiliations:** Louis Calder Center - Biological Field Station, Fordham University, Armonk, NY, USA; Department of Biological Sciences and Center for Urban Ecology, Fordham University, Bronx, NY, USA

**Keywords:** priority effects, community assembly, historical contingency, *Batrachochytrium dendrobatidis*, amphibian, microbiome

## Abstract

Priority effects in host-associated microbiomes can influence not only community composition and structure, but also community functions, such as disease resistance. However, evidence for these priority effects remains scarce. Past studies suggest that amphibian protection from chytridiomycosis, caused by the pathogenic fungus *Batrachochytrium dendrobatidis* (Bd), is related to antifungal bacterial composition on host skin. Priority effects in these bacterial communities may influence susceptibility to Bd, but this possibility has not been tested. Using *in vitro* microcosms, we demonstrated that priority effects can influence interactions among amphibian-associated microbes. We observed strong priority effects irrespective of high antifungal ability such that the Bd-inhibitory potential of two strongly inhibitory bacterial species did not always produce higher levels of Bd-inhibition. This result suggests that interactions may be more complex than previously thought. Additionally, our results suggest that priority effects between commensal and pathogenic taxa can be either facilitatory or inhibitory, with the strength and direction of this effect dependent on the composition of the community. Thus, changes in assembly may lead to varying levels of Bd infection, influencing how we might augment amphibian-associated microbiomes to conserve taxa currently at risk of extinction.

## Introduction

Many bacteria, archaea, and microeukaryotes form symbioses with plant and animal hosts acting as an extension of the host’s innate immune system (Cho and Blaser 2012, Cani 2018). Researchers have taken advantage of the natural competition that exists among microbes to develop probiotic or prebiotic treatments that favor putatively beneficial microbes over pathogens. While the number of studies examining multi-species communities has increased to reflect the growing knowledge of the complexity of host microbiomes, replicating these results in the host have shown mixed success (Sánchez et al. 2013, Papadimitriou et al. 2015, Akhter et al. 2015). This is likely due in part to our limited knowledge on microbe-microbe and microbe-environment (itself a dynamic, living organism) interactions, or more specifically, how these come together historically to influence community function (Lawley et al. 2012, Fraune et al. 2015). Thus, successful probiotic development and application requires deeper ecological understanding of microbiome assembly, stability, and variability.

Generally, the assembly of a community results from species dispersal from the regional species pool to the local community filtered by interactions among species and the environment as well as chance fluctuations in demography, environment, and dispersal (Gilpin and Hanski 1991, Wilson 1992). Slight changes in the order of species arrival can result in a historical contingency which can interact with deterministic assembly mechanisms (e.g., niche) to create lasting effects on local community structure and function (Palmgren 1926, Fukami 2015). For instance, a species may exhibit an absolute fitness advantage during assembly allowing them to dominate in the community simply by being the first colonizer. This phenomenon is known as a priority effect (Drake 1991), and similar to competitive lottery models, it offers a framework for exploring the role of both deterministic and stochastic factors in assembly (Verster and Borenstein 2018). Despite the increasing number of studies acknowledging the presence of assembly rules over the last century, few general principles have emerged that allow one to predict when history matters and when it does not (Lawton 1999, Fukami and Morin 2003, Fukami 2015). This challenge is often due to the fact that obtaining relevant historical information for diverse ecological communities is difficult or impossible leading studies to infer the outcome of assembly based on pairwise species co-occurrences. Thus, few studies have explored how the actual process of community formation (e.g., the initial sequence of colonization) provides context for which assembly mechanisms are most important in each system.

The amphibian-pathogen relationship presents a relevant conservation system for testing the presence of priority effects in host-associated microbial communities and how they lead to changes in community function. Scientists have found that protection from chytridiomycosis (caused by the invasive fungal pathogen, *Batrachochytrium dendrobatidis* (Bd)) is directly related to native bacterial composition (i.e., the types of bacterial species and their relative abundances) on host skin (Harris et al. 2006, Becker and Harris 2010). Here, cutaneous composition is determined by both the host’s ability to meet the resource requirements of microbial species and the ability of those species to impact the environment on the host (Kueneman et al. 2014). Additionally, skin is a highly exposed environment (as compared to internal host environments), so community composition is largely impacted by stochastic events that result in random loss or acquisition microbes. In various animals, passive dispersal and drift are often more important than selection in skin microbiome assembly (Adair et al. 2018, Chiarello et al. 2019, Tong et al. 2019, Barnes et al. 2020b). These stochastic processes are major drivers of variation within and among hosts, which in addition to host-specific attributes likely explain why individuals within a population can vary in their susceptibility to particular pathogens. Thus, while disease is driven by pathogen invasion, host susceptibility is greatly influenced by microbial dysbiosis wherein the abundance of beneficial taxa is highly regulated by assembly from the source community (Verster and Borenstein 2018, Wei et al. 2019, Wilber et al. 2020).

Furthermore, bacteria and Bd likely overlap in resource use (Barr 1969, Piotrowski et al. 2004), and both have been found to modify their environment by producing and releasing metabolites (Brucker et al. 2008b, 2008a, Loudon et al. 2014, Rollins-Smith et al. 2015). These factors have been shown to result in strong priority effects in other disease systems where stochastic differences in microbiome composition resulted in altered community function (Sharon et al. 2013, Lam and Monack 2014, Devevey et al. 2015). For this reason, elucidating the conditions affecting beneficial bacterial establishment onto amphibian host skin will improve our ability to manipulate these communities for enhanced disease protection.

This study aims to explore how both historically contingent and deterministic assembly processes affect (1) the abundance of Bd-inhibitory bacterial taxa in synthetic communities, and (2) how these processes may lead to varying levels of Bd infection *in vitro*. We acknowledge that our *in vitro* microcosms are less taxonomically complex than natural amphibian host microbiomes. However, by first reducing the complexity associated with feedbacks between a living host and its microbiome, we were able to focus on the microbe-microbe dynamics that could either directly or indirectly influence pathogen abundance. Over the course of 7 d, we tracked microbial dynamics between common amphibian microbes and Bd in microcosms that varied in the order of microbial introduction. We hypothesized that the first colonizers would preempt niche space resulting in priority effects. Additionally, we also hypothesized that the strength of priority effects between bacteria and Bd would depend on the identity of bacterial colonists with stronger priority effects produced among species with higher resource use overlap or those that are capable of niche modification (Vannette and Fukami 2014, Shaani et al. 2018, Estrela et al. 2020). For example, we predicted that *Janthinobacterium lividum* and *Stenotrophomonas rhizophila* (both highly anti-fungal Proteobacteria) would have an increased influence on each other’s abundances as a result of overlapping functional roles (Scheuring and Yu 2012). Overall, the results of this study will improve our understanding of the complex role of stochastic and deterministic assembly mechanisms in Bd infection enhancing our ability to address conservation crises.

## Materials and Methods

### Study organisms

We selected three bacterial species isolated from the skin of a woodland salamander, *Plethodon cinereus*, in New York, USA in 2017 (Barnes et al. 2020a). These species varied in their ability to inhibit *Batrachochytrium dendrobatidis* (Bd) from the total bacterial community collected: two strongly Bd-inhibitory (*Janthinobacterium lividum* and *Stenotrophomonas rhizophila*) and one weakly Bd-inhibitory species (*Bacillus cereus*). These species are also commonly found on amphibian skin regardless of host species (Woodhams et al. 2017, Wolz et al. 2018, Barnes et al. 2020a) and could be easily isolated and cultured, making them relevant to addressing amphibian disease susceptibility (Bletz et al. 2013). The Bd-inhibitory ability of each bacterial species was confirmed by in-house challenge assays with Bd (strain JEL 423, University of Maine, Orono, USA) and with the Antifungal Isolates Database (Woodhams et al. 2015, Becker et al. 2015).

### Experimental design

We used a 3 x 3 x 4 factorial design with three orders of Bd introduction, three combinations of bacterial species, and four timepoints (**Figure 1**). Bd-introduction consisted of: i) simultaneous introduction of Bd and bacteria, and ii) sequential introduction, where either bacteria or Bd were introduced 2 d before the other. All pairwise bacterial combinations were included in the experiment.

**Figure 1.**
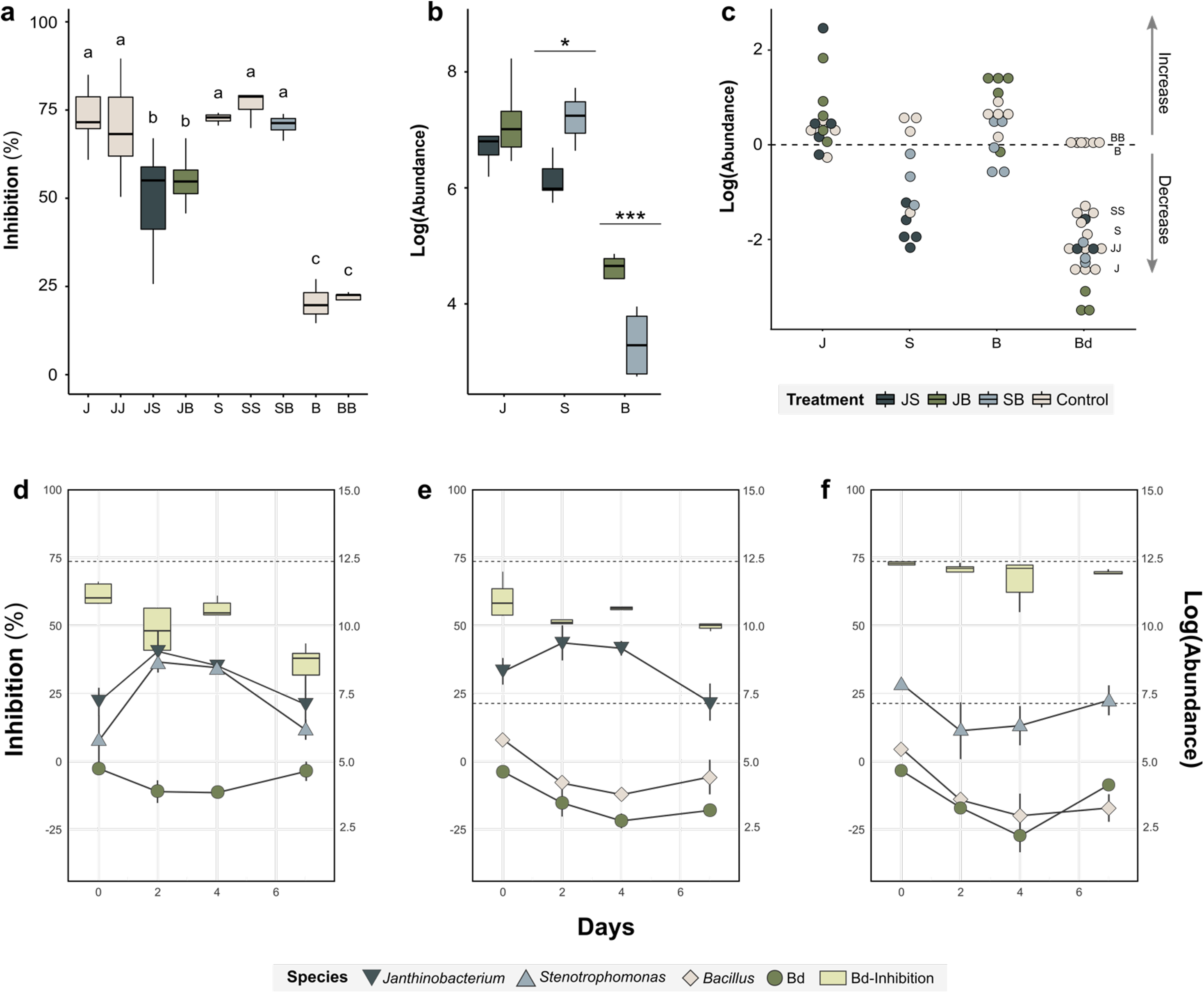
Simultaneous introduction of bacteria and Bd after 7 d (*n* = 5). **(a)** Bd-inhibition for each bacterial combination. **(b)** Bacterial population abundance shown by isolate where color denotes bacterial combination. **(c)** Normalized difference in microbial population abundance between treatments and control (e.g., J, S, and B with Bd, and Bd-alone) for each bacterial species and Bd at 7 d. Values of zero indicate no difference in population as compared to the control. **(d-f)** Microbial population abundance and Bd-inhibition in microcosms over time. Bd-inhibition is on the left y-axis and population abundance is on the right y-axis. Bd-inhibition by 96-well CFS-challenge assay, and population abundance was measured via qPCR. From **(d-f)** bacterial combinations are: *Janthinobacterium x Stenotrophomonas, Janthinobacterium x Bacillus, and Stenotrophomonas x Bacillus*. Bacteria indicated as: J = *Janthinobacterium*, S = *Stenotrophomonas*, B = *Bacillus*, where individual letters indicate controls (e.g., J = 100 ul, JJ = 200ul or 2X volume). Dotted lines indicate mean Bd-inhibition of each individual bacterial species with Bd. Boxplots show median and interquartile range. Statistical significance between groups shown as letters for **(a)** (*p* < 0.05) and stars for **(b)** (* denotes *p* < 0.05 and *** denote *p* < 0.001).

From our long-term culture collection, each bacterial isolate was diluted to 1:10^4^ cells and grown on 1% tryptone agar at 23°C for 24 h to produce colonies. A 1 cm^2^ agar plug (as colony size varied by species) was then placed in 1% tryptone broth at 23°C for 24 h to create our experimental stock cultures. To produce our Bd stock culture, 1 ml of Bd was plated on 1% tryptone agar and grown at 23°C for 4 d after which we decanted and transferred the zoospores to new 1% tryptone broth and grew these liquid cultures at 23°C for 48 h. Microcosms consisted of 2 ml microcentrifuge tubes containing 700 ul of 1% tryptone broth, 100 ul Bd (~2.0 x 10^6^ zoospores measured via hemocytometer), and 100 ul of each bacterial species in pairs. Positive controls included microcosms with bacterial isolates by themselves (in 1X and 2X volumes) with Bd. Each treatment contained five independent replicates per time point.

### Population abundance and inhibition estimation

Immediately following each sampling time point, replicate samples were centrifuged for 5 min at 10,000g to pellet microbial cells. The supernatant was removed and filtered through a 0.22-um cellulose acetate syringe filter to isolate the cell-free supernatant (CFS). DNA was extracted from pelleted cells using PrepMan™ Ultra Sample Preparation Reagent (Applied Biosystems, Foster City, CA) following manufacturer’s protocols. DNA extracts were then cleaned using Serapure beads to remove cell debris and PCR inhibitors.

Estimation of microbial population abundance was conducted using quantitative PCR with genera-specific primers (**Table S1**). Bacterial primers were adapted from previous studies that identified gene sequences unique at or below genus-level. Multiple sequences for the target genes, *vio*, *ggpS*, and *motB*, as well as available whole genomes of the microbes of interest were downloaded from NCBI and aligned in MUSCLE using ClustalW. All primers were tested against the four microbes of interest and an additional 15 isolates from our culture collection to confirm specificity. Quantitative PCR was performed on an Applied Biosystem QuantStudio™ 3 machine with SYBR® Green reagents. PCR reactions were run in 20 ul-volume: 10 ul of PowerUp™ SYBR Green Master Mix (2X), 1 ul each of genera-specific forward and reverse primers (500 nM), and 4 ul of DNA. Amplification consisted of initial UDG activation for 2 min at 50°C and denaturation for 2 min at 95°C followed by 40 cycles of: 15 s at 95°C, 15 s at primer-specific annealing temperature (**Table S1**), and 1 min at 72°C. After amplification, a melting step was performed under default conditions. Each bacterial stock culture was grown on 1% tryptone agar at 23°C until colonies could be counted to estimate colony forming units (CFU) for bacterial abundance standard curves.

Bd-inhibition was measured using the 96-well method developed by Bell et al. (2013). In brief, the CFS of each microcosm was assayed with Bd (50 ul CFS: 50 ul Bd-zoospore suspension) in duplicate and grown for 10 days at 23°C. Optical density was measured on an Infinite 200 Pro microplate reader (Tecan Trading AG, Männedorf, Switzerland) at 0, 4, 7, and 10 d. Percent Bd-inhibition was then calculated from the absorbance readings following procedures outlined in Barnes et al. (2020).

### Statistical Analyses

Nested ANOVAs were performed using a mixed-effect model with bacterial combination, introduction order, and time as fixed effects. Post-hoc comparisons on least-mean squares were performed with a Tukey adjustment and Bray-Curtis dissimilarities between communities were visualized via NMDS. The strength of priority effects of Bd on bacteria and bacteria on Bd was quantified by adapting the methods in Vannette & Fukami (2014). Strength was calculated as the log of the ratio between the abundance of species *i*, here measured by qPCR, when it was introduced after species *j* versus before species *j* (*P_ij_* = ln [*A*(*i*)_*ji*_ / *A*(*i*)_*ij*_], where subscripts indicate introduction order). To adjust for the length of time in the microcosm between our ordered introduction treatments, we compared abundances at 7 d for “after” treatments to abundances at 4 d in “before” treatments. Homogeneity of variance was confirmed using Levene’s test and two-way ANOVAs with Tukey HSD post-hoc comparisons were used to compare priority effects across treatments. All analyses and plots were completed using the *vegan*, *nlme*, *lsmeans*, and *ggplot2* packages in R version 3.6.1 (Oksanen et al. 2013, Wickham 2016, Pinheiro et al. 2017, Lenth and Lenth 2018, R Core Team 2019).

## Results

### Microbe-microbe interactions under simultaneous introductions

Of the three bacterial combinations tested, we found that none of the combinations worked additively or synergistically with respect to Bd-inhibition (**Figure 1a**). In all cases, Bd-inhibition was lower than expected as compared to the additive effect of each bacterial species alone with Bd (*F* = 73.9, *p* < 0.001). As expected, we found no significant difference in inhibition between the two highly Bd-inhibitory bacteria, *Janthinobacterium* (J) and *Stenotrophomonas* (S; mean_J_ = 72.8 ± 7.7%, mean_S_ = 72.1 ± 6.3%), as well as no significant difference between J or S controls and JJ or SS controls which were inoculated with 2X the volume of bacteria (*p*_J-JJ_ = 0.95, *p*_S-SS_ = 0.96). Additionally, Bd-inhibition in control microcosms of the weakly inhibitory bacteria, *Bacillus* (B), was significantly different from Bd-inhibition in highly inhibitory J or S control microcosms (*p*_J-B_ < 0.001, *p*_S-B_ < 0.001). In microcosms containing more than one bacterial species (i.e., bacterial combination treatments), we found that SB microcosms resulted in Bd-inhibition 1.4X and 1.3X higher than that found in JS (*p*_JS-SB_ < 0.001) and JB microcosms (*p*_JB-SB_ < 0.001). There was no significant difference in Bd-inhibition between JS or JB microcosms (*p*_JS-JB_ = 0.72).

When we compared the abundance of bacteria and Bd (**Figure 1b-c**), JS microcosms had the highest average bacterial abundance. While we found no significant difference in the abundance of *Janthinobacterium* in any of our microcosms (including treatments and controls), JS and JB had slightly higher *Janthinobacterium* abundance compared to J or JJ controls. In contrast, *Stenotrophomonas* abundance decreased in both JS and SB treatments compared to S and SS controls but was 1.2X higher in SB microcosms compared to JS microcosms (*p*_JS-SB_ = 0.02). SB microcosms also had significantly higher Bd-inhibition at 7 d compared to all other bacterial combination treatments (*p* < 0.05, JS < JB < SB; **Figure 1d-f**). Despite this, JB microcosms showed the largest decrease in Bd abundance compared to all other microcosms including J and S controls (**Figure 1c**; *F* = 122.5, *p* < 0.001).

### Simulating Bd infection with ordered introductions

When we compared simultaneous and ordered introduction treatments, we found a significant effect of introduction order and bacterial combination on Bd abundance and Bd-inhibition (**Table 1a-b**). Additionally, we found significant differences in the abundances of: *Janthinobacterium* with time, *Stenotrophomonas* with bacterial combination, and *Bacillus* with both introduction order and bacterial combination (**Table 1c-e**).

**Table 1.**
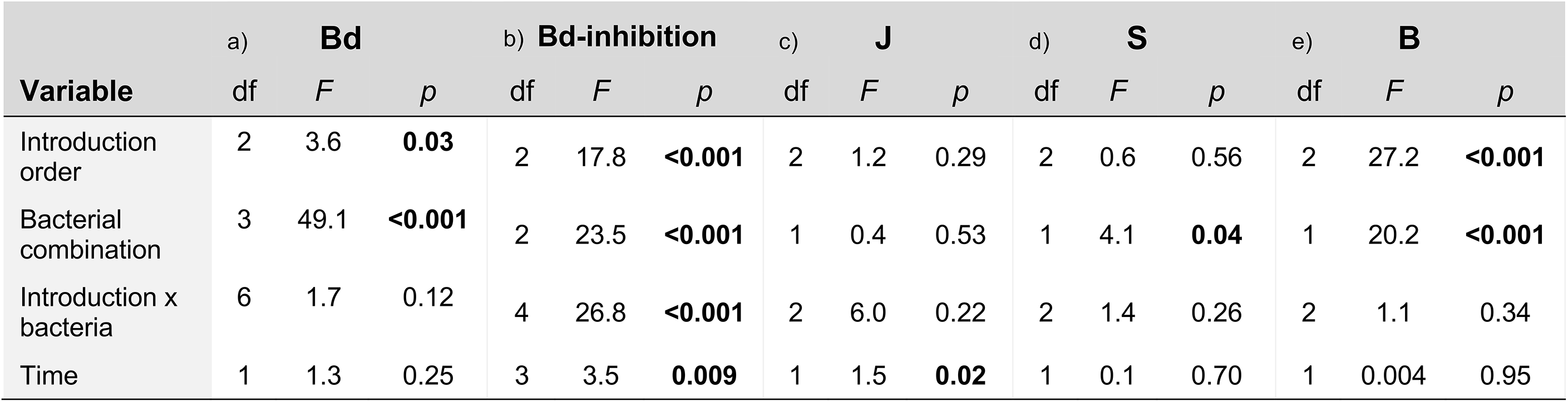
Results from two-way ANOVA relating a) Bd abundance, b) Bd-inhibition, c) *Janthinobacterium* (J) abundance, d) *Stenotrophomonas* (S) abundance, and e) *Bacillus* (B) abundance throughout the experiment to introduction order (Bd before, Bd after, or simultaneous introduction), bacterial combination (JS, JB, or SB), their interaction, and time of sampling (0, 2, 4, or 7 days). *P*-values < 0.05 shown in bold.

While all ordered introductions of Bd and bacteria resulted in significantly less Bd-inhibition than simultaneous introductions at 7 d (**Figure 2**; *F* = 11.8, *p* < 0.001), they varied in their effect on microbial abundance. In microcosms containing both putatively highly Bd-inhibitory bacteria, *Janthinobacterium* and *Stenotrophomonas* (JS), we found no significant difference in *Janthinobacterium* or *Stenotrophomonas* abundance between either ordered treatment at 7 d (**Figure 2a**; *F*_J_ = 2.9, *p* = 0.10; *F*_S_ = 0.3, *p* = 0.72). However, both species had a slower growth rate in bacteria-first microcosms (noted as JS/Bd), which corresponded with significantly lower Bd-inhibition at 4 d as compared to Bd-first or simultaneous microcosms (**Figure 2d-e**; *F* = 9.2, *p* = 0.009). When *Janthinobacterium* was paired with the putatively weakly Bd-inhibitory bacteria, *Bacillus* (JB), we found a significant increase in Bd abundance in ordered treatments compared to simultaneous treatments (**Figure 2b**; grouping of both ordered treatments to the left of NMDS axis 1 and simultaneous treatments to the right). Additionally, both Bd-inhibition and *Janthinobacterium* abundance varied significantly with time in JB ordered treatments (**Figure 2f-g**; *p_Bd-inhibition_* < 0.01, *p_J_* = 0.02). Despite an increase in *Janthinobacterium* abundance prior to 4 d, Bd-inhibition significantly decreased between 2 and 4 d in both ordered treatments (*p_BacteriaFirst_* = 0.007; *p_BdFirst_* = 0.004) resulting in 0.3-0.5X inhibition relative to JB simultaneous introduction and J or JJ control treatments.

**Figure 2.**
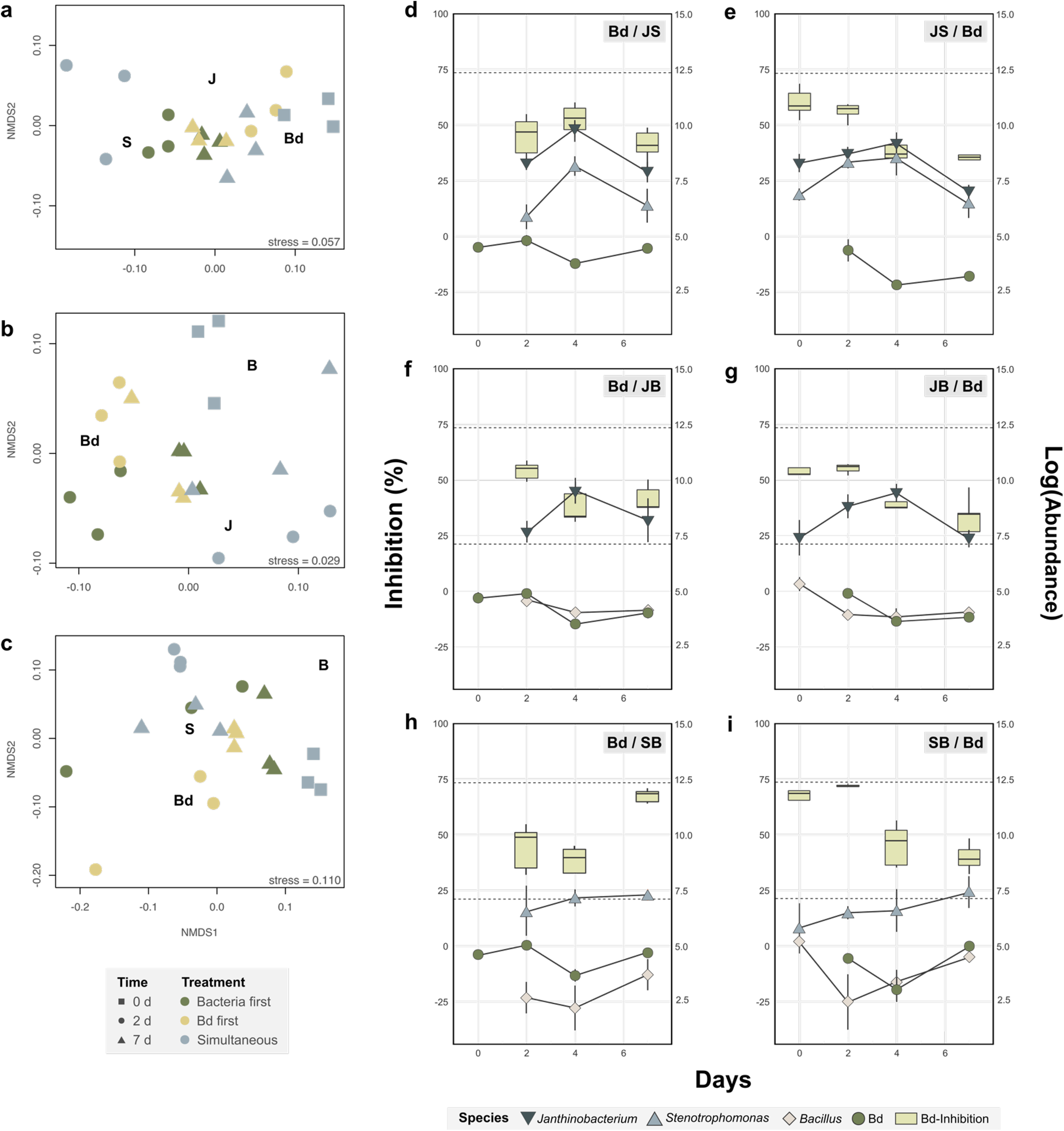
Microbial community composition of microcosms. **(a-c)** NMDS plots of microbial community microcosms (*n* = 3) under each bacterial combination (listed as JS, JB, BC) on each day of introduction (0 d and 2 d) and on the final day of the experiment (7 d). **(d-i)** Microbial population abundance and Bd-inhibition in ordered introduction microcosms (*n* = 5) over time shown as **(d,f,h)** introduction of Bd before bacteria, and **(e,g,i)** introduction of bacteria before Bd. Bd-inhibition is on the left y-axis and population abundance is on the right y-axis. Bd-inhibition by 96-well CFS-challenge assay, and population abundance was measured via qPCR. Bacteria indicated as: J = *Janthinobacterium*, S = *Stenotrophomonas*, B = *Bacillus*, where individual letters indicate controls (e.g., J = 100 ul, JJ = 200ul or 2X volume). Dotted lines indicate mean Bd-inhibition of each individual bacterial species with Bd. Boxplots show median and interquartile range.

In microcosms containing *Bacillus* and *Stenotrophomonas* (SB), we found that Bd-inhibition at 7 d was 2X higher when bacteria were introduced after Bd than before (*p* < 0.001; no significant difference between SB_BdFirst_, SB_Sim_, or J and S controls at 7 d). Despite high abundance of *Stenotrophomonas* and *Bacillus* when bacteria were introduced before Bd (**Figure 2c**), Bd-inhibition decreased and Bd abundance increased with time (**Figure 2h-i**).

### Priority effects between bacteria and Bd

To quantify the effects of introduction order on the microbes associated with amphibian skin, we compared the abundance of both bacteria and Bd in bacteria-first and Bd-first treatments. When Bd was the first colonizer, we found significant priority effects on each bacterial isolate, but effects varied in direction and strength based on bacterial combination (*p*_Bacteria_ < 0.001, *p*_Culture_ = 0.002, *p*_Bacteria:Culture_ = 0.004). In all combinations, Bd had an inhibitory effect on *Janthinobacterium* and a facilitatory effect on *Bacillus* (**Figure 3a**; *p*_J-B_ < 0.001, *p*_JS-JB_ = 0.99, *p*_JB-SB_= 0.91). Unlike *Janthinobacterium* and *Bacillus*, the strength and direction of Bd’s priority effect on *Stenotrophomonas* varied significantly with bacterial combination. When *Stenotrophomonas* was co-cultured with *Janthinobacterium*, Bd had an inhibitory effect, and when co-cultured with *Bacillus*, Bd had a facilitatory effect on *Stenotrophomonas* abundance (*F*_JS-SB_ = 24.9, *p*_JS-SB_ = 0.002). Notably, the strength of the inhibitory effect of Bd on *Stenotrophomonas* was 3X greater than the facilitatory effect. When bacteria were the first colonizers, they had an inhibitory priority effect on Bd except for in the SB microcosms (**Figure 3b**; *F* = 22.3, *p* < 0.001) where the growth rate of Bd at 7 d was higher when bacteria were introduced first (**Figure 2i**).

**Figure 3.**
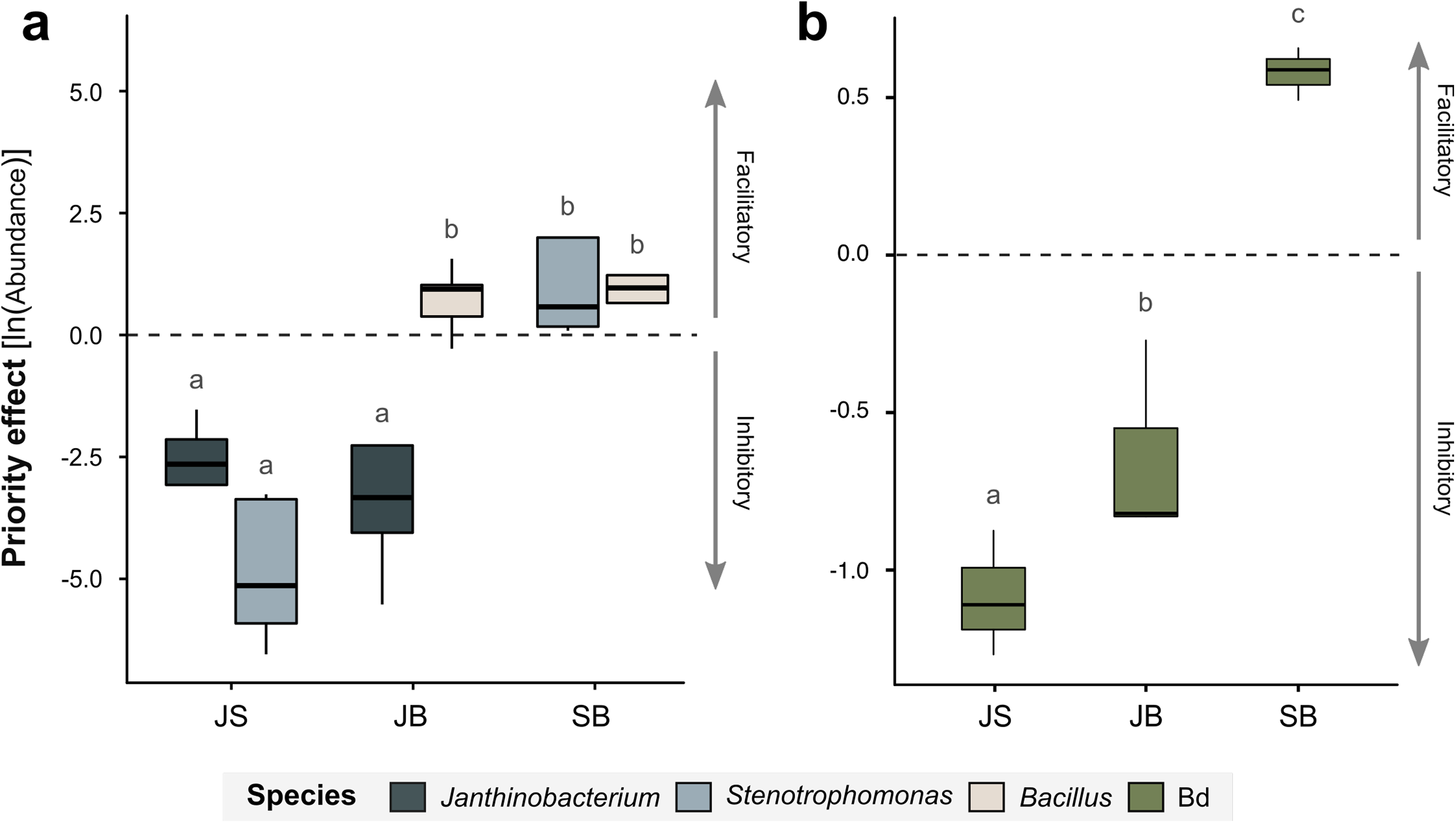
Priority effects in ordered introduction microcosms with different bacterial combinations (*n* = 5) where **(a)** effect of Bd on bacterial abundance, and **(b)** effect of bacteria on Bd abundance. Strength of the priority effect was calculated as the natural logarithm of the ratio of species abundance under “before vs. after” ordered introduction. Positive values indicate facilitatory effects and negative values indicate inhibitory effects. Bacterial combinations indicated by color and as: J = *Janthinobacterium*, S = *Stenotrophomonas*, B = *Bacillus*. Boxplots show median and interquartile range with outliers (outside 1.5X the interquartile range) shown as points. Statistical significance between groups (calculated separately for **a** and **b**) shown as letters (*p* < 0.05).

## Discussion

This study used an *in vitro* model of the amphibian microbiome to explore how changes in community assembly (e.g., species introduction order and composition of community members) influence Bd infection. We found that all treatments using combinations of the two highly Bd-inhibitory bacteria, *Janthinobacterium lividum* and *Stenotrophomonas rhizophila*, resulted in lower Bd-inhibition than either microbe alone. Further, we observed strong priority effects between all bacterial taxa and Bd irrespective of potential antifungal ability, but that the direction of Bd’s priority effect (either facilitatory or inhibitory) on each bacterial taxon varied with isolate identity and, in the case of *S. rhizophila*, the identities of the other bacteria it was cultured with. These results highlight the complexity of interactions that can occur during even simple microbial assembly—the dynamics of which are often overlooked among putatively commensal and pathogenic host-associated microbes.

### Insights from in vitro assembly experiments

The use of simplified, *in vitro* microbial communities was critical to uncovering and characterizing potential priority effects in this system. Much of the research currently underway regarding the use of bacteria for mitigating chytridiomycoses in wild amphibian populations uses *in vitro* co-culture microcosms to identify candidate microbes which have since been applied directly to live hosts (Bletz et al. 2013, Woodhams et al. 2014, Rebollar et al. 2016). We know that the amphibian microbiome is influenced by host-associated (e.g., behavior, developmental stage, immunity), abiotic (e.g., pH, salinity, temperature), and biotic factors (e.g., pathogen presence, food resources, environmental reservoirs; Rebollar et al. 2016). Further, many of these factors are interconnected, which inherently has led to a trade-off in these studies: maintaining complexity of the host-associated microbiome or gaining specific, mechanistic insights into microbial ecology and assembly. This, at least in part, explains why *in vivo* experimental bioaugmentation trials have had mixed results. Thus, we decided to return to the *in vitro* microcosm-based experiments long-used in assembly theory (Woodruff 1912, Gause 1934, Drake et al. 1996, Jessup et al. 2004) because they allowed us to control for both host-associated and abiotic factors to reveal the range of possible commensal-pathogen interactions that can influence higher level organization and/or function. The goal is then to apply the insights gained from these simplified models to systems with increasing complexity and live hosts.

### Single vs. co-culture probiotic additions

Prior to manipulating microbiome introduction order, we established a baseline for Bd-inhibitory function and microbial abundances in single- and co-culture microcosms using three common amphibian bacteria. We found that microcosms were often less inhibitory to Bd when composed of two bacterial species. In particular, co-culturing of two putatively Bd-inhibitory bacteria, *Janthinobacterium lividum* and *Stenotrophomonas rhizophila*, resulted in significantly lower Bd-inhibition than either bacteria cultured alone with Bd. Co-cultures containing *Bacillus cereus* showed significantly higher Bd-inhibition than *B. cereus* alone, but Bd-inhibition in these microcosms was still significantly lower than the Bd-inhibitory potential of *J. lividum* or *S. rhizophila* by themselves. While two-species bacterial communities are not nearly as complex as actual amphibian microbiomes, this work reveals that more species-rich bacterial mixtures may not always result in higher disease prevention—likely due to the complexity of interactions among microbial species, including antagonism and facilitation (Reid et al. 2011, Gerritsen et al. 2011, Loudon et al. 2016). Competition between microbes should be assessed not only between Bd and beneficial bacteria, but also between bacteria within the probiotic and those common to the host’s natural environment (Woodhams et al. 2014, Kelsic et al. 2015). Given this study and others, Bd resistance is likely not a single trait, but is due to a combination of factors: less niche space, an increase in abundance and specificity of antimicrobial metabolites, and indirect competition—many of which are not often predictable based on phylogenetic relatedness (Belden and Harris 2007, Becker et al. 2015, Barnes et al. 2020a).

### Priority effects influence pathogen load

Additionally, stochastic processes (i.e., dispersal and drift) are known to occur alongside deterministic processes to drive variation in communities, especially at the microbial scale (Belyea and Lancaster 1999, Chase 2003, Nemergut et al. 2013, Bannar-Martin et al. 2018). By manipulating introduction order of beneficial, Bd-inhibitory microbes, we found that both stochastic (e.g., introduction order) and deterministic processes (e.g., microbial interactions) contribute to a community’s resistance or resilience to pathogen invasion. As hypothesized, we found that Bd-inhibitory bacteria (JS) often exert inhibitory priority effects on Bd when introduced first likely due their advantage in being the first to take up space and resources. JB microcosms were the only other microcosms to reach comparable decreases in Bd abundance, which is surprising for two reasons: 1) this microcosm contained one highly and one weakly Bd-inhibitory bacteria, and 2) simultaneous introduction of bacteria and Bd goes against the current practice of applying beneficial bacteria to sterilized amphibian skin (i.e., prior to Bd exposure).

When Bd was introduced first (Bd-first microcosms), it exerted a mixture of positive and negative priority effects depending on bacterial identity. The presence of weak facilitatory priority effects from both bacteria and Bd in SB microcosms may explain why Bd-inhibition dropped and Bd abundance increased despite both bacteria also increasing in abundance at 7 d in bacteria-first treatments. In particular, bacteria-first SB microcosms had the greatest increase in *Bacillus* abundance and also the lowest Bd-inhibition, despite no significant change in *Stenotrophomonas* abundance. Persistence in a community can be influenced by not only competition, but also syntrophic partnerships and nutrient cross-feeding (Seth and Taga 2014). These interactions might explain why we saw a disconnect between Bd abundance and Bd-inhibition measured by our CFS assay at several timepoints. These findings were unexpected, and we encourage more work to be done to tease apart the mechanisms driving these interactions at the cost or benefit of the pathogen.

Finally, our analysis revealed the importance of time on microbial abundance in synthetic microcosms. We believe that the decrease in Bd-inhibition at 4 d in all bacterial combinations containing *Janthinobacterium* (JS and JB) was reflective of the decrease of both J and Bd abundance and thus a potential decrease in Bd-inhibitory metabolites. However, decreases in microbial abundance with time could also have been due to the confounding effect of not providing any supplemental media or colonists during the experiment. Still, the decrease in *Janthinobacterium* abundance in co-culture treatments was greater than in J or JJ controls, suggesting that there could be additional factors associated with co-culturing affecting *Janthinobacterium* growth and function. One such factor could be that both *J. lividum* and *S. rhizophila* occupy similar niche space, as both are phylogenetically related (soil Proteobacteria) and have similar growth requirements. Shared niche space combined with the competitive effect of Bd could act to prevent either bacteria from dominating and instead create coexistence in the community. This would be beneficial with regard to long-term maintenance of Bd-inhibitory function on amphibian skin and should be explored beyond our 7-day timescale. Additionally, the main antifungal metabolite produced by *J. lividum*, violacein, has also been shown to impact Gram-positive bacteria (Choi et al. 2015). This might explain why JB microcosms (containing the Gram-positive bacteria, *Bacillus*) showed lower *B. cereus* abundance than some treatments containing the bacterial combination SB.

## Conclusion

In sum, this research has shown that bacteria do not always act synergistically to inhibit Bd. Among these four amphibian microbes, we show that interactions and resulting function can be historically contingent and result in priority effects. Thus, stochastic variation in microbiome assembly may give an advantage to either commensal or pathogenic microbial colonizers, which can ultimately lead to varying levels of disease susceptibility among hosts. These potential long-lasting effects associated with colonization order and species composition in pathogen systems is rarely explored, but as shown here, can lead to significant differences in community function. Finally, our results suggest that development of amphibian probiotic treatments will be most successful if we look beyond microbial identity, continue to characterize communities based on function, and consider both stochastic and deterministic assembly processes as important drivers of disease outcomes.

---

**Table S1.**
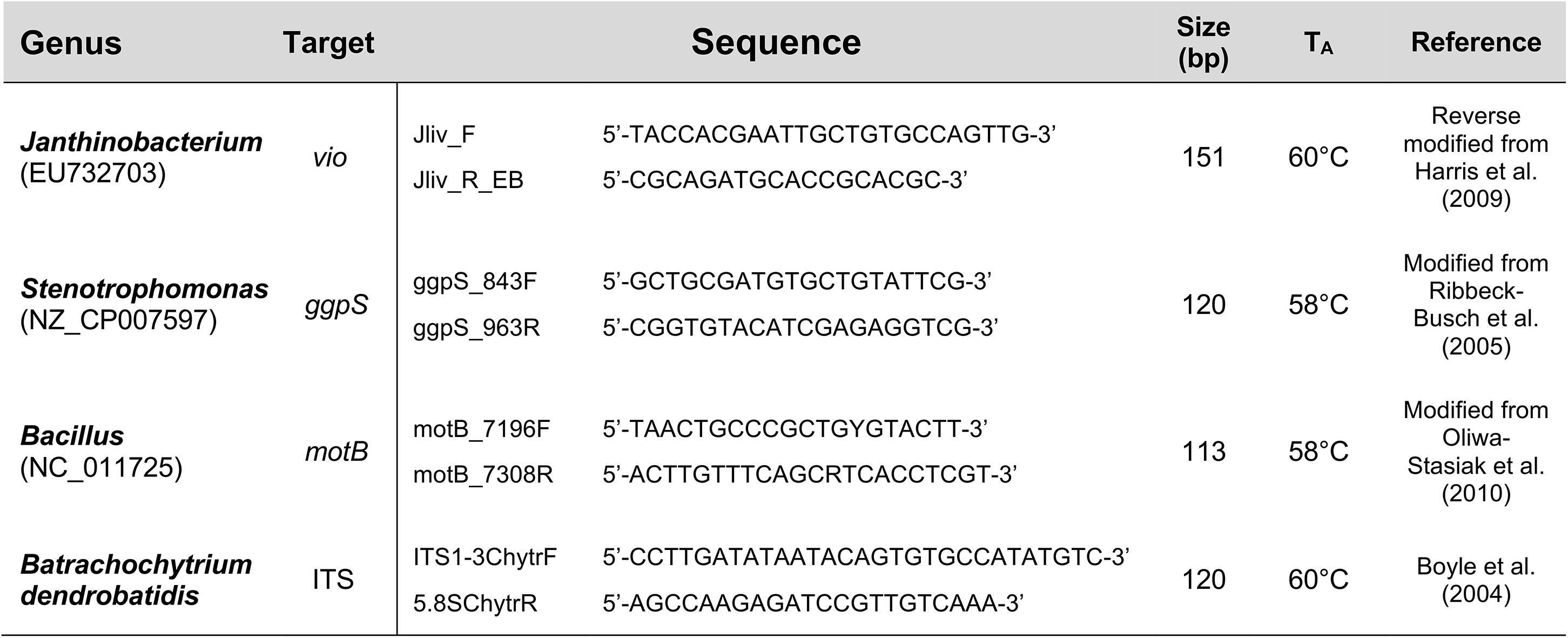
PCR primers used for identification and quantification of bacteria and Bd during qPCR. GenBank ascension numbers for target genes are provided below each bacterial genus.

